# DeepNano-blitz: A Fast Base Caller for MinION Nanopore Sequencers

**DOI:** 10.1101/2020.02.11.944223

**Authors:** Vladimír Boža, Peter Perešíni, Broňa Brejová, Tomáš Vinař

## Abstract

**Motivation:** Oxford Nanopore MinION is a portable DNA sequencer that is marketed as a device that can be deployed anywhere. Current base callers, however, require a powerful GPU to analyze data produced by MinION in real time, which hampers field applications.

**Results:** We have developed a fast base caller DeepNano-blitz that can analyze stream from up to two MinION runs in real time using a common laptop CPU (i7-7700HQ), with no GPU requirements. The base caller settings allow trading accuracy for speed and the results can be used for real time run monitoring (i.e. sample composition, barcode balance, species identification, etc.) or pre-filtering of results for more detailed analysis (i.e. filtering out human DNA from human–pathogen runs).

**Availability and Implementation:** DeepNano-blitz has been developed and tested on Linux and is available under MIT license at https://github.com/fmfi-compbio/deepnano-blitz

## 1 Introduction

We introduce DeepNano-blitz, a very fast base caller for Oxford Nanopore MinION sequencers. MinION is a pocket-sized DNA sequencing machine which measures electric current as a DNA molecule passes through a nanopore. These electric signals can be then translated into sequences using a base caller software (such as Guppy [Wick et al., 2019] or Chiron [Teng et al., 2018]) with error rate of about 10-15%. MinIONs are extremely portable, and therefore suitable for experiments in the field or deployment in a clinical setting; in fact, all equipment required to extract DNA, prepare sequencing libraries, and perform the sequencing fits in a regular travel suitcase [Edwards et al., 2016].

Unfortunately, in a struggle to increase the accuracy, the new versions of base callers have become slower over time. To keep up with approx. 1.5-2.0 million signal readouts per second coming from a successful MinION run, one currently needs a powerful GPU with compute capability at least 6.1. Such requirements are often not met by regular laptop computers, which are in other ways perfectly capable of running a MinION. Heavy GPU usage also implies high energy consumption, which hampers field deployment of MinION sequencing.

Here, we introduce a new base caller DeepNano-blitz, which can keep up with one or even two MinION sequencers on a single i7-7700HQ 4-core laptop CPU with no GPU, at the cost of slightly reduced accuracy. Our intended use is real-time monitoring of a sequencing run (such as to ascertain proportions of barcodes in a sample, examine the ratio of human vs. pathogen DNA, select a minority of interesting reads for in-depth analysis, etc.), even though some of the data may need to be re-analyzed later with a more accurate tool.

## 2 Methods

DeepNano-blitz is based on a bi-directional recurrent neural network (similarly to Guppy [Wick et al., 2019], Chiron [Teng et al., 2018], or DeepNano [Boža et al., 2017]) which is heavily optimized for performance (see Supplementary Figure S1). We start by taking a median-normalized raw signal as an input and fork it into two channels (identity and squared input). The first part of the network preprocesses this input by temporal convolution, followed by temporal max-pooling with stride 3, and tanh layer. The main part of the network consists of four GRU layers in alternating directions. As the last step, we use a softmax layer, each position predicting one of -, A, C, G, T. The network is trained using CTC loss [Graves et al., 2006]. In decoding, we use a beam search with tunable beam size, dropping beams if the last step probability is less than a tunable threshold. The details of training and testing sets used are shown in Supplementary Section S1.

To achieve greater speed, DeepNano-blitz is written in Rust and hand-optimized. We employ cache-aware memory layouts, fast approximations of sigmoid/tanh functions [Mineiro, 2011], and Intel MKL library for matrix multiplication. Our current implementation manages to run over 15 floating point multiplications per CPU cycle which is close to the architectural maximum.

## 3 Results

To evaluate the speed and accuracy of our tool, we have used a benchmark data set of R9.4.1 *Klebsiella pneumoniae* [Wick et al., 2019]. The base calls were mapped to the reference using minimap2 [Li, 2016] and the read accuracy is computed as one minus the ratio of the alignment edit distance and the length of the base call. We report the median read accuracy. Figure 1 shows the comparison of various settings of the DeepNano-blitz against fast and high-accuracy settings of Guppy 3.4.4(cpu version). In the fastest mode, the DeepNano-blitz runs over 100x faster than Guppy high accuracy and approx. 13x faster than Guppy in fast mode. The accuracy difference between the version that can keep up with 2 million readouts per second (width64-beam5) and Guppy in fast mode is less than 2 percentage points. For more complete results, including a human data set [Jain et al., 2018], see Supplementary Table S2).

**Figure 1:**
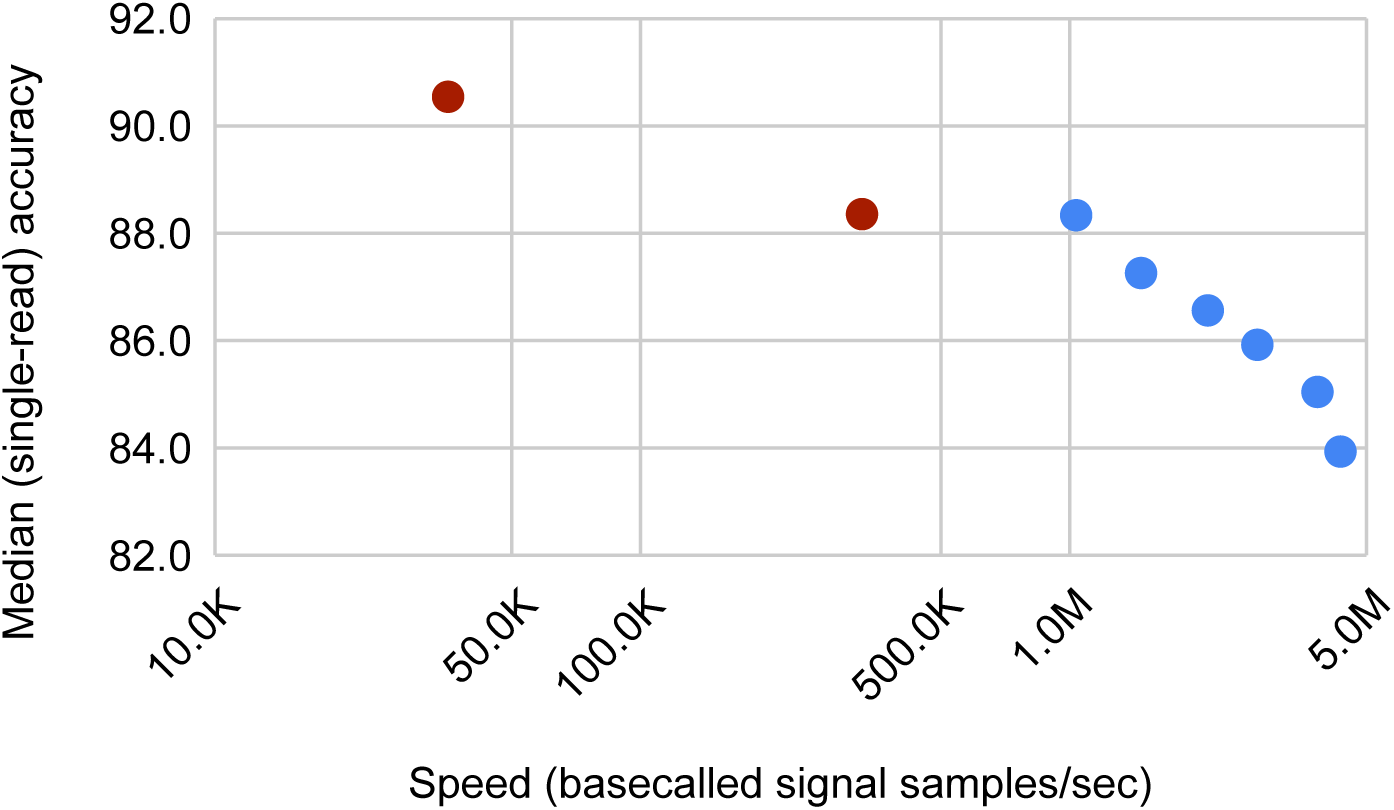
Comparison of accuracy versus speed of basecalling. Red: Guppy (leftmost point corresponds to high-accuracy setting), Blue: DeepNano-blitz (using different settings of neural network size to adjust the speed-accuracy trade-off). A single MinION can produce up to 1.5-2M signal samples per second. Note that the x-axis is in log scale.

It is true that a few percentage points in read accuracy can sometimes significantly impact the accuracy of downstream analysis. To ascertain this effect, we have analyzed ZymoBIOMICS Microbial Community Standards data set [Oxford Nanopore Technologies, 2019], [Nicholls et al., 2019] with both DeepNano-blitz and Guppy 3.4.4. Each read was base called and using minimap2, it was mapped to the reference genomes to estimate the composition of the sample. Guppy in high accuracy mode resulted in 94.5% reads successfully mapped, while the fastest version of DeepNano-blitz mapped 90.7% of reads. There were no significant differences between the estimated proportions of reads (Supplementary Table S3).

We have also ascertained the ability of DeepNano-blitz to monitor the balance of a barcoded sample, using a public data set of 12 barcoded bacterial samples [Wick et al., 2018]. After base calling, we have used guppy barcoder with standard settings to classify reads into barcodes. As expected, DeepNano-blitz results in lower recall (approx. 26% of reads classified as unknown barcode with width64-beam5 setting vs. 17% with guppy fast mode); yet, there is no significant difference in precision (approx. 96% with width64-beam5 vs. 97% with guppy fast mode) and there are no significant differences in estimates of barcode composition of the sample between DeepNano-blitz and Guppy 3.4.4(Supplementary Table S4). Note that guppy barcoder is optimized for errors typically made by guppy and with custom barcode recognition settings, recall of DeepNano-blitz may be improved.

## 4 Discussion

DeepNano-blitz provides a fast alternative to Oxford Nanopore base callers for analysis of MinION data. It allows trading accuracy for speed and enables real-time data analysis without requirement of a powerful GPU. We believe, that DeepNano-blitz enhances the ability to deploy MinION sequencing in the field and enables building custom analysis pipelines monitoring MinION sequencing runs in real time.

## Acknowledgements

This research was supported in part by funding from the European Union’s Horizon 2020 research and innovation programme under grant agreement No 872539, and by grants from the Slovak Research and Development Agency (APVV-18-0239) and VEGA (1/0458/18 to TV and 1/0463/20 to BB).

