## Supplementary Material for "DeepNano-blitz: A Fast Base Caller for MinION Nanopore Sequencers"

#### S1 Neural Network, Training and Testing Sets

The topology of our neural network is outlined in Figure S1.

We have used the mixture of the following public data sets to train the models:

- *Taiyaki set:* Data set of 50k (downsampled to 5k) R9.4.1 reads from a PCR amplified DNA of *E. coli* (SCS110), *H. sapiens* (NA12878), and *S. cerevisiae* (NCYC1052), published by Oxford Nanopore as a part of Taiyaki software (https://github.com/nanoporetech/taiyaki/).
- *E. coli set:* A sample of 2804 ultra-long native R9.4.1 reads of *E. coli* (MG1655) from Loman Lab (https://lab.loman.net/2017/03/09/ultrareads-for-nanopore/).
- *Human training:* A sample of reads from chr1 and chr2 of native R9.4.1 reads (flowcell FAB49164) from human reference standard CEPH1463 from nanopore whole human genome sequencing project [Jain et al., 2018].

For testing, the following data sets were used:

- *Klebsiella*: a benchmark set of native R9.4 *K. pneumoniae* reads [Wick et al., 2019]
- *Human testing:* a sample of native R9.4.1 reads from chr14, chr15, chr16 (flowcell FAB49164) from human reference standard CEPH1463 from nanopore whole human genome sequencing project [Jain et al., 2018]
- *Zymoset test:* Oxford Nanopore Technologies Zymo Mock Community data on R9.4.1 (https://github.com/nanoporetech data)
- *Barcoding test:* A sample of 4000 reads from the data set of 12 barcoded bacterial samples PRJEB28450 [Wick et al., 2018]

The basic characteristics of each data set is shown in Table S1.

*First two authors contributed equally to the study.

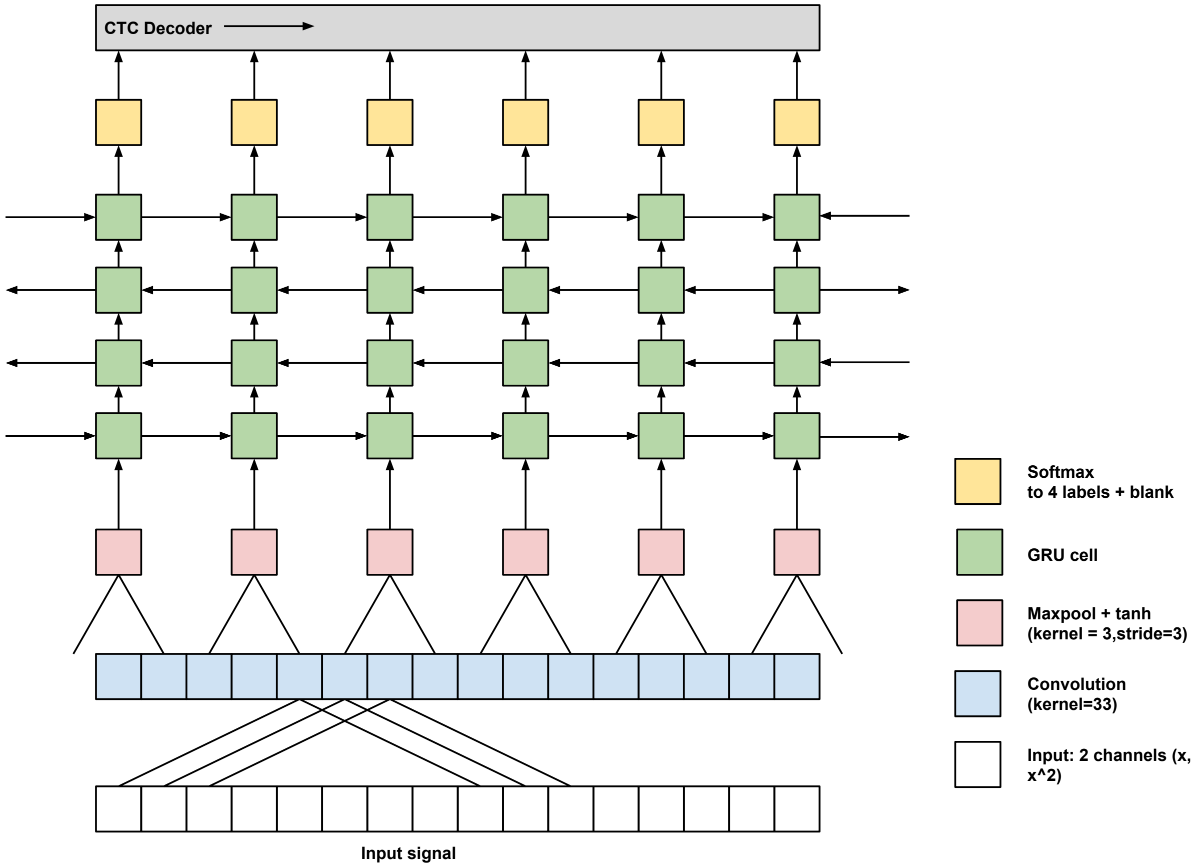


Figure S1: Topology of the neural network.

| Data Set | # reads | Total length (in bp) | Mean read length (in bp) | Median read length (in bp) |
|---|---|---|---|---|
| Taiyaki set | 5000 | 28.8 Mbp | 5769 | 5459 |
| E. coli training set | 2804 | 90.6 Mbp | 32319 | 21758 |
| Human training set | 1323 | 11.5 Mbp | 8705 | 7111 |
| Human testing set | 305 | 2.7 Mbp | 8936 | 6231 |
| Klebsiella | 5000 | 105.8 Mbp | 21667 | 16119 |
| Zymoset | 4000 | 110.2 Mbp | 2755 | 2752 |
| Barcoding test | 4000 | 44.0 Mbp | 11003 | 6847 |

Table S1: Overview of training and testing sets used in the study.

### S2 Evaluation and Performance

| Base caller and parameters | Signal readouts processed per second | Klebsiella pneumoniae | | Human | |
|---|---|---|---|---|---|
| | | % mapped | Median accuracy | % mapped | Median accuracy |
| width 48, beam 1 | 4.4M | 98.6 | 84.0 | 84.3 | 80.6 |
| width 48, beam5 | 3.8M | 98.8 | 85.1 | 84.9 | 81.6 |
| width 56, beam5 | 2.8M | 98.9 | 85.9 | 86.1 | 82.3 |
| width 64, beam5 | 2.1M | 99.0 | 86.6 | 85.5 | 83.4 |
| width 80, beam5 | 1.5M | 99.0 | 87.3 | 86.1 | 84.3 |
| width 96, beam5 | 1.0M | 99.3 | 88.4 | 87.4 | 85.9 |
| guppy 3.4.4 fast | 328.6K | 99.5 | 88.4 | 89.1 | 85.1 |
| guppy 3.4.4 hac | 35.1K | 99.5 | 90.6 | 89.6 | 87.4 |

Table S2: *Speed and accuracy at various settings.* Evaluation of base calling at various settings trading accuracy for speed. The tests were performed on a laptop with 4-core i7-7700HQ CPU. We report the median read accuracy over all reads in the corresponding testing set.

| Base caller and parameters | Mapped % | *B. subtilis* | *C. neoformans* | *E. coli* | *E. faecalis* | *L. fermentum* | *L. monocytogenes* | *P. aeruginosa* | *S. aureus* | *S. cerevisiae* | *S. eneterica* |
|---|---|---|---|---|---|---|---|---|---|---|---|
| width 48, beam 1 | 90.7 | 15.19% | 1.43% | 14.32% | 14.57% | 5.43% | 16.05% | 4.68% | 13.97% | 1.64% | 12.73% |
| width 48, beam 5 | 91.0 | 15.22% | 1.46% | 14.37% | 14.52% | 5.44% | 16.02% | 4.69% | 13.91% | 1.63% | 12.75% |
| width 56, beam 5 | 91.3 | 15.22% | 1.42% | 14.34% | 14.56% | 5.43% | 16.05% | 4.70% | 13.93% | 1.61% | 12.75% |
| width 64, beam 5 | 91.5 | 15.23% | 1.43% | 14.35% | 14.56% | 5.43% | 16.03% | 4.69% | 13.89% | 1.63% | 12.76% |
| width 80, beam 5 | 91.7 | 15.27% | 1.41% | 14.37% | 14.57% | 5.42% | 16.01% | 4.69% | 13.91% | 1.63% | 12.73% |
| width 96, beam 5 | 92.0 | 15.22% | 1.41% | 14.35% | 14.56% | 5.42% | 16.10% | 4.69% | 13.92% | 1.62% | 12.71% |
| guppy 3.4.4 fast | 94.3 | 15.26% | 1.45% | 14.36% | 14.60% | 5.46% | 16.07% | 4.68% | 13.79% | 1.64% | 12.69% |
| guppy 3.4.4 hac | 94.5 | 15.26% | 1.45% | 14.32% | 14.63% | 5.50% | 16.05% | 4.66% | 13.80% | 1.65% | 12.67% |

Table S3: *Comparison of metagenomic composition estimates.* There are no significant differences between the estimates of the sample composition based on guppy and DeepNano-blitz base calls.

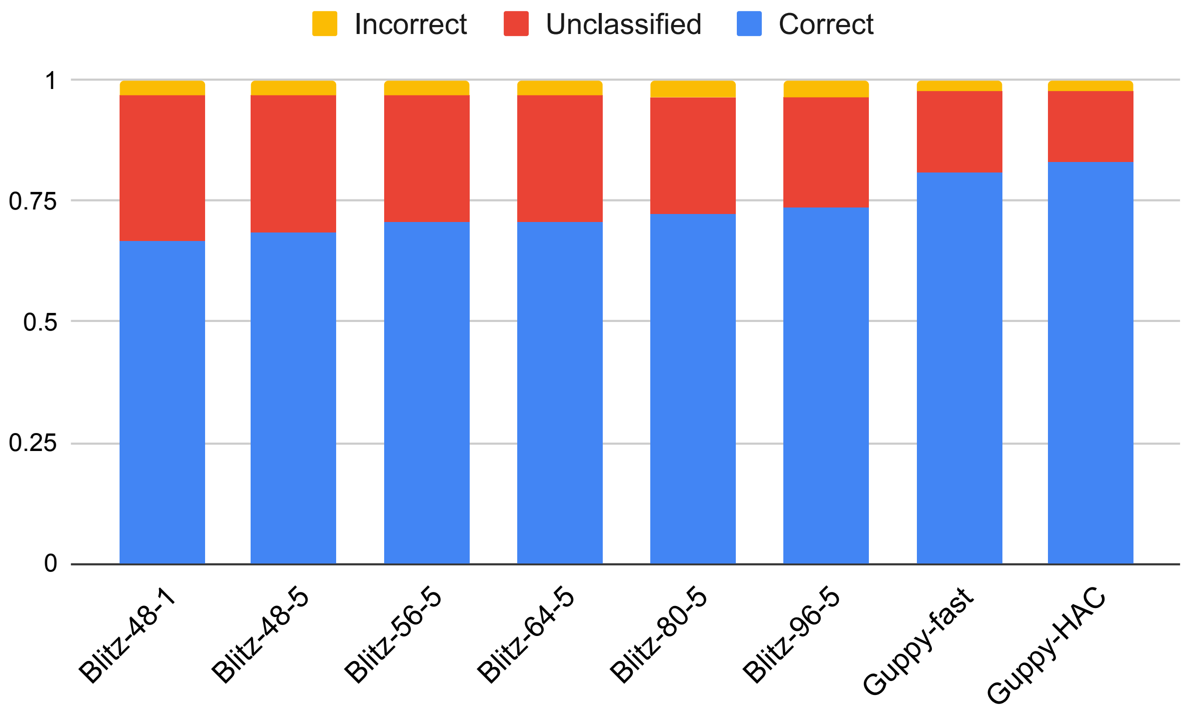


Figure S2: Recall and precision of barcode classification.

| **Base caller** | **classified %** | **Identified barcode** | | | | | | | | | | |
|---|---|---|---|---|---|---|---|---|---|---|---|---|
| | | **01** | **02** | **03** | **04** | **05** | **06** | **07** | **08** | **09** | **11** | **12** |
| width 48, beam 1 | 63.70% | 4.59% | 12.52% | 7.61% | 9.26% | 8.99% | 20.05% | 5.73% | 2.28% | 16.01% | 9.73% | 3.22% |
| width 48, beam5 | 65.42% | 4.47% | 12.23% | 7.57% | 9.32% | 9.09% | 20.06% | 5.73% | 2.25% | 16.28% | 9.71% | 3.29% |
| width 56, beam5 | 67.67% | 4.51% | 12.08% | 7.68% | 9.49% | 9.01% | 19.43% | 5.43% | 2.44% | 17.21% | 9.49% | 3.21% |
| width 64, beam5 | 67.33% | 4.49% | 12.14% | 7.72% | 9.91% | 8.95% | 19.49% | 5.68% | 2.78% | 16.12% | 9.65% | 3.04% |
| width 80, beam5 | 69.47% | 4.39% | 12.56% | 7.63% | 9.79% | 8.92% | 19.36% | 5.54% | 2.45% | 16.70% | 9.50% | 3.17% |
| width 96, beam5 | 70.80% | 4.27% | 12.46% | 7.77% | 10.10% | 8.90% | 19.67% | 5.51% | 2.65% | 16.28% | 9.22% | 3.18% |
| guppy 3.4.4 fast | 76.75% | 4.20% | 12.70% | 7.43% | 10.23% | 8.89% | 19.51% | 5.28% | 2.83% | 16.42% | 9.28% | 3.22% |
| guppy 3.4.4 hac | 78.78% | 4.19% | 12.66% | 7.36% | 10.66% | 8.85% | 19.07% | 5.27% | 2.76% | 16.72% | 9.20% | 3.24% |

Table S4: Barcode balance estimation.
